# Progesterone modulates theta oscillations in the frontal-parietal network

**DOI:** 10.1101/2020.01.16.909374

**Authors:** Justin Riddle, Sangtae Ahn, Trevor McPherson, Susan Girdler, Flavio Frohlich

## Abstract

The neuroactive metabolites of the steroid hormones progesterone (P4) and testosterone (T) are GABAergic modulators that influence cognitive control, yet the specific effect of P4 and T on brain network activity remains poorly understood. Here, we investigated if a fundamental oscillatory network activity pattern related to cognitive control, frontal midline theta (FMT) oscillations, are modulated by steroids hormones, P4 and T. We measured the concentration P4 and T using salivary enzyme immunoassay and FMT oscillations using high-density electroencephalography (EEG) during the eyes-open resting state in fifty-five healthy female and male participants. Electrical brain activity was analyzed using Morlet wavelet convolution, beamformer source localization, background noise spectral fitting, and phase amplitude coupling analysis. Steroid hormone concentrations and biological sex were used as predictors for scalp and source-estimated theta oscillations and for top-down theta-gamma phase amplitude coupling. Elevated concentrations of P4 predicted increased FMT oscillatory amplitude across both sexes, and no relationship was found with T. The positive correlation with P4 was specific to the frontal-midline electrodes and survived correction for the background noise of the brain. Using source localization, FMT oscillations were localized to the frontal-parietal network. Additionally, theta amplitude within the frontal-parietal network, but not the default mode network, positively correlated with P4 concentration. Finally, P4 concentration correlated with increased coupling between FMT phase and posterior gamma amplitude. Our results suggest that P4 concentration modulates brain activity via upregulation of theta oscillations in the frontal-parietal network and increased top-down control over posterior cortical sites.

**Significance Statement:** The neuroactive metabolites of the steroid hormones progesterone (P4) and testosterone (T) are GABAergic modulators that influence cognitive control, yet the specific effect of P4 and T on brain network activity remains poorly understood. Here, we investigated if a fundamental oscillatory network activity pattern related to cognitive control, frontal midline theta (FMT) oscillations, are modulated by steroids hormones, P4 and T. Our results suggest that P4 concentration modulates brain activity via upregulation of theta oscillations in the frontal-parietal network and increased top-down control over posterior cortical sites.

## Introduction

The metabolites of the sex steroids, progesterone (P4) and testosterone (T), are allosteric modulators of the GABA_A_ receptor and impact cognition (Zurkovsky et al. 2007; Schumacher et al. 2014; Celec, Ostatníková, and Hodosy 2015), as well as other biological and affective systems. The progestogens (e.g. P4) and androgens (e.g. T) are classes of sex steroids that differentially engage neural activity patterns in functional neuroimaging (Peper et al. 2011) and differentially influence cognition (Pletzer, Petasis, and Cahill 2014). In particular, a recent study found that elevated concentrations of progesterone (P4) correlate with greater performance in the N-back, working memory task (Hidalgo-Lopez and Pletzer 2017) suggesting a role for P4 in boosting cognitive control signals. Consistent with a role of P4 in cognitive control, increased resting state connectivity within the frontal-parietal network in functional magnetic resonance imaging positively correlated with P4 concentrations (Syan et al. 2017). Despite emerging evidence for a role of P4 in cognitive control, there has yet to be an investigation into a possible connection between P4 concentrations and the prominent cognitive control signal in electrical brain activity: frontal midline theta (FMT) oscillations (4-7Hz) (see (Cavanagh and Frank 2014) for review).

T concentrations are associated with alterations in social cognition: social decision-making (Eisenegger and Naef 2011), aggression (Montoya et al. 2012), and social status (Eisenegger, Haushofer, and Fehr 2011). Consistent with the role of T in social cognition, greater concentrations of T are correlated with decreased functional connectivity between the orbito-frontal cortex (critical region for social cognition) and the amygdala (critical region for processing emotionally salient perception) (Ackermann et al. 2012). Frontal theta oscillations also have been shown to play a role in social cognition (Tendler and Wagner 2015; Narayanan et al. 2011). Therefore, T concentration may also be predictive of FMT oscillations.

We investigated the relationship between the steroid hormones, P4 and T, and FMT oscillations using salivary enzyme immunoassay and recorded high-density electroencephalography (EEG) during the resting state in men and women. Using source localization, we investigated the neural origin of FMT oscillations and correlated theta activity in the frontal-parietal network to steroid hormone concentrations. Finally, we hypothesized that the mechanism by which steroid hormones contribute to cognitive control is via coupling of the phase of prefrontal theta oscillations to the amplitude of gamma activity (30-50 Hz) in parietal-occipital cortex.

## Materials and Methods

Data from fifty-five participants was pooled across three experiments (22 from NCT0324450 (Sheffield et al. 2019), 14 from NCT03243084 (Ahn et al. 2019), and 19 from NCT03178344. In each of these studies, we sampled steroid hormone concentrations via saliva sample and collected resting-state electrical brain activity via high-density EEG. All experiments were approved by the institutional review board at the University of North Carolina at Chapel Hill and data were collected at the Carolina Center for Neurostimulation. Participants were screened prior to enrollment with the following inclusion criteria: no personal or immediate family history of neurological or psychiatric illness, no ongoing psychotherapy treatment or use of medication to treat a neurological or psychiatric illness, no major head injury or brain surgery, no brain implants (including cochlear implants), no history of cardiovascular disease. All participants were required to pass a urinary drug test and female participants were screened with a pregnancy test. Participants were asked to maintain a regular sleep schedule prior to the experimental sessions, as well as abstain from alcohol and caffeine for the 24 hours prior to the visit.

### Steroid Hormone Enzyme Immunoassay

Steroid hormone concentrations were assessed via four saliva samples taken throughout a single day: before breakfast, before lunch, before dinner and before bed. Participants were instructed not to eat for an hour before giving the sample. The data were collected within a week of the EEG and sent to Labrix for assay (Clackamas, OR, USA). Labrix pooled the four samples into a single sample estimate and conducted enzyme immunoassays for P4 and T. The P4 (4-pregenen-3,20-dione) enzyme immunoassay kit was provided by Salimetrics Inc (State College, PA, USA) with a standard curve ranged from 6.1-600.0 pg/mL, a coefficient of variation for intra-assay precision from 4.0-8.4% and inter-assay precision of 5.5-9.6%, and a strong correlation with serum P4 (r(35) = 0.80). The kit measured free P4 in the saliva that is not bound to serum proteins and is considered biologically active. Cross-reactivity was greatest for corticosterone (0.192%), 17 α-hydroxyprogesterone (0.0723%), 11-deoxycortisol (0.0195%) and cortisone (0.0106%). There was no detectable (<0.004%) cross-reactivity with T, estradiol, DHEA, or cortisol (Salimetrics Salivary Progesterone Enzyme Immunoassay Kit Item No. 1-2502). The T enzyme immunoassay kit was the Pantex Salivary Direct Testosterone EIA Kit by PANTEX Division of Bio-Analysis (Santa Monica, CA, USA) with a standard curve ranged from 10.0-2400 pg/mL, a coefficient of variation for intra-assay precision from 2.1-5.1% and inter-assay from 3.1-6.4%. 1-2% of T is free in plasma, and this concentration is highly correlated with free T in saliva. Like P4, the kit measures only biologically active T. Cross-reactivity was greatest for 5 α-dihydro-testosterone (5.47%), 5 α-androstone-3 β, 17 β-diol (2.75%), methyl testosterone (1.60%), and 11 α-OH testosterone (1.03%). There was low cross-reactivity with P4 (0.28%), 17 β-estradiol (0.173%), DHEA (0.003), and cortisol (0.002%) (Pantex Salivary Direct Testosterone EIA Kit Catalog No. 635). These kits show low cross-reactivity between our steroids of interest. The immunoassays have an exponential distribution. Therefore, raw values were log-transformed prior to all statistical analysis.

### High Density Electroencephalography

EEG data were collected with a high-density 128-channel electrode net at 1000 Hz (HydroCel Geodesic Sensor Net) and EGI system (NetAmps 410, Electrical Geodesics Inc., OR, USA). The impedance of each electrode was ensured to be below 50 kΩ at the start of each session. Four-minutes (five-minutes in 19 participants from NCT03178344) of eyes-open resting-state EEG was collected.

The resting-state EEG data were preprocessed using custom scripts in MatLab and the EEGLAB toolbox (Delorme and Makeig 2004). All EEG data were downsampled to 250Hz with anti-aliasing filtering and band-pass filtered from 1 to 50 Hz. The data were then preprocessed using an artifact subspace reconstruction algorithm to remove high-variance and reconstruct missing data (Mullen et al. 2013). Channels that were found in the previous step to contain above threshold noise were interpolated. All data was common average referenced. An infomax independent component analysis (ICA) was performed to separate plausible neural activity from eye blinks, eye movement, muscle activity, heartbeats, and channel noise (Jung et al. 2000). All ICA components were visually inspected and components corresponding to noise were manually rejected. Resting-state EEG data were epoched into two second windows for scrubbing. Each epoch was visually inspected in time domain, and epochs with abnormal spikes were rejected from analysis.

Our analysis was restricted to the 90 channels on the scalp, referred to as data channels. The fast Fourier transform was applied to each two-second epoch and the average spectral amplitude was calculated. To estimate frontal midline theta (FMT), we first calculated the average amplitude from 4 to 7 Hz, the canonical theta band, for each data channel. Then, these data were log-transformed and normalized for each participant using the z-transformation across the 90 data channels. Finally, FMT was then defined as the resulting spatially-normalized spectral amplitude in electrode FCz.

### Experimental Design and Statistical Analysis

We used an independent Student’s t-test between sex for each steroid hormone concentration and FMT oscillatory amplitude to test for any differences in our metrics driven by sex. We were interested in the contribution of steroid hormone concentrations and biological sex to FMT. Therefore, we ran a correlation between each of our independent variables, P4 and T, with our dependent variable, FMT. To confirm the topological specificity of a significant correlation with FMT, we correlated theta amplitude from every data channel with the independent variable. We hypothesized that a genuine correlation with FMT would be confined to a cluster surrounding FCz.

### Correction for Aperiodic Signal

Recent evidence suggests that the aperiodic signal in the background of the brain may play a meaningful role in modulating cognition and predicting healthy aging (Voytek et al. 2015). As the aperiodic signal of the brain, a 1/f power distribution, steepens there will be an increase in theta and delta power (2-7Hz). The change in slope could lead to the erroneous conclusion that theta oscillations are increasing, when the more parsimonious explanation is a steepening of the 1/f slope (Gao, Peterson, and Voytek 2017). A true oscillation under this framework is defined as a band-limited increase in spectral power over and above the 1/f background spectra (He 2014). Therefore, we calculated the 1/f background spectra and removed this trend from our data to confirm that the measured theta oscillations represent true oscillations. We calculated the 1/f background spectra as a linear fit to the log(amplitude) and log(frequency) of each participant. As the alpha oscillation is prominent in EEG during the resting state, we calculated the aperiodic signal as the linear fit to amplitude values from 2-4Hz and 40-50Hz (see (Voytek et al. 2015) for a similar method). After fitting the data for each participant, we subtracted the fit amplitude across all frequencies in order to calculate 1/f corrected amplitude. We then correlated the peak corrected theta amplitude values with steroid hormone concentrations.

### Network Analysis

Our large sample size and use of a 128-channel EEG system provided a methodological foundation for source localization analysis to investigate the spatial origination of FMT oscillations. Source localization was run using the Dynamical Imaging of Coherent Sources (DICS) beamformer algorithm implemented in the FieldTrip toolbox (Oostenveld et al. 2011). For each two second epoch of resting-state data, we calculated the spectral amplitude for each channel via 5-cycle Morlet wavelet convolution at 6 Hertz. Next, we calculated the cross-spectral density matrix to estimate the phase difference and shared amplitude of each channel pair for source localization. The beamformer algorithm was run using a lead field calculated in the Montreal Neurological Institute (MNI) space using standard skull, skin, and brain tissue estimates provided in the FieldTrip toolbox. Theta amplitude was source localized into MNI space for each two second epoch and then averaged across epochs. The theta amplitude estimates from source localization were not normally distributed. Therefore, the data were log-transformed. We applied spatial normalization using the z-transformation for all voxels within a grey-matter mask in MNI space derived from tissue segmentation estimates provided by the SPM12 toolbox (Penny et al. 2011). For display purposes, an independent t-test between participants with high and low FMT amplitude in sensor-space (median split) was run on theta-amplitude for every voxel in source-space.

Investigation into the temporal and spatial engagement of the human brain during cognitive control has found two primary signals: one spatial and one temporal. Electrophysiology researchers find increased FMT oscillations and hypothesized that these signals originate from the anterior cingulate cortex or medial prefrontal cortex directly beneath scalp sensors with peak FMT amplitude (see (Cavanagh and Frank 2014) for review), whereas functional neuroimaging researchers find increased functional connectivity and activation within the FPN as a function of increased cognitive control as driven by task demands (see (Badre and Nee 2018) for review). When dipoles in homologous brain regions are aligned in opposite directions (e.g. medial to lateral), the resulting electric field is found in the space between the two dipoles where the field inverts. This principle of electric field modeling is found in task-evoked electrical activity. For example, in auditory tasks, peak auditory-evoked electrophysiological signal is found in central midline electrode, Cz; but via source localization auditory activity is found to be driven by bilateral auditory cortex (Stropahl et al. 2018). Therefore, we hypothesized that source localization of FMT oscillations would reveal greatest theta amplitude in bilateral frontal cortex (Sasaki et al. 1994; Sasaki et al. 1996) and not in superior frontal gyrus or medial prefrontal cortex.

Bilateral middle frontal gyri have network membership in the task-positive frontal-parietal network (FPN), whereas anterior superior frontal gyrus and medial prefrontal cortex have network membership in the default mode network (DMN) (Yeo et al. 2011). To quantify the degree to which theta oscillations were source localized to the FPN as opposed to the DMN, we averaged theta amplitude in grey matter voxels of the FPN and DMN. FPN and DMN were defined as the 6^th^ and 7^th^ network from the Yeo atlas derived from resting-state functional connectivity analysis (Yeo et al. 2011). This network analysis provides a critical dimensionality reduction by which 10,000s of voxels are reduced to two networks; and, therefore, addresses the multiple comparisons problem. Analyses were run using custom code written in MatLab using functions from SPM12 and Fieldtrip.

### Phase Amplitude Coupling

Phase amplitude coupling (PAC) is a proposed mechanism for top-down cognitive control in the human brain, in which the phase of low frequency oscillations in the prefrontal cortex are coupled to high frequency oscillations in posterior regions of the brain (see (Canolty and Knight 2010) for review). We hypothesized that the phase of FMT might play a role in directing high frequency gamma oscillations [30-50Hz] in parietal-occipital cortex (Berger et al. 2019). Therefore, we calculated phase amplitude coupling between theta phase in FCz and gamma amplitude in parietal-occipital electrodes. We ran the Hilbert transform on the time series of each electrode to extract theta phase and gamma amplitude. Next, we calculated phase amplitude coupling between the theta phase (θ) of FCz and the gamma amplitude (M) of each parietal-occipital electrode across all time points 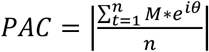. We generated a null distribution of 1000 PAC values by randomly shifting the gamma amplitude time series by a minimum of 10% relative to the theta phase time series. Finally, PAC_Z_ is the z-normalized genuine PAC value from the permutation-generated null distribution for each electrode pair for each participant (Cohen 2014). PAC can only be positive. Therefore, we ran one-tailed statistics to test for the presence of PAC against the null hypothesis that PAC_Z_ is not different from 0.

### Code accessibility

Scripts used for these analyses can be found in our code repository for this study on the Open Science Framework (https://osf.io/6j253/).

## Results

Biological sex difference for steroid hormone concentrations and FMT oscillations are displayed in Table 1. We found that females had lower T concentrations compared to males. There was no significant difference in age, P4, or FMT between females and males. P4 was positively correlated with FMT oscillations for all participants (r(54) = 0.366, p = 0.0061) (Figure 1A). The positive relationship between P4 and FMT was consistent for females (r(22) = 0.439, p = 0.036) and males (r(32) = 0.328, p = 0.067), albeit less so in males. T was not correlated with FMT oscillations in either sex (r(54) = 0.107, p = 0.435) (Figure 1B).

**Table 1.** Biological sex differences. Descriptive estimates, mean and standard deviation (SD), for all independent and dependent variables are displayed by sex. Data were normalized prior to t-test according to the description in Methods. *** p < 0.001.

| MEAN (SD) | Females<br>(N = 23) | Males<br>(N = 32) | Females & Males<br>(N = 55) | Sex Difference (f - m) |  |
| --- | --- | --- | --- | --- | --- |
|  |  |  |  | t(53) | d |
| Age [y] | 31.1 (13.7) | 27.2 (12.3) | 28.8 (12.9) | 1.10 | .30 |
| Progesterone (P4) [pg/mL] | 142.3 (103.7) | 112.4 (76.9) | 124.9 (89.5) | 0.16 | .04 |
| Testosterone (T) [pg/mL] | 20.2 (8.51) | 94.3 (38.5) | 63.3 (47.3) | **-13.4\*\*\*** | 3.62 |
| Frontal Midline Theta [μV] | 0.994 (0.31) | 0.928 (0.24) | 0.956 (0.27) | 0.13 | .04 |

**Figure 1.**
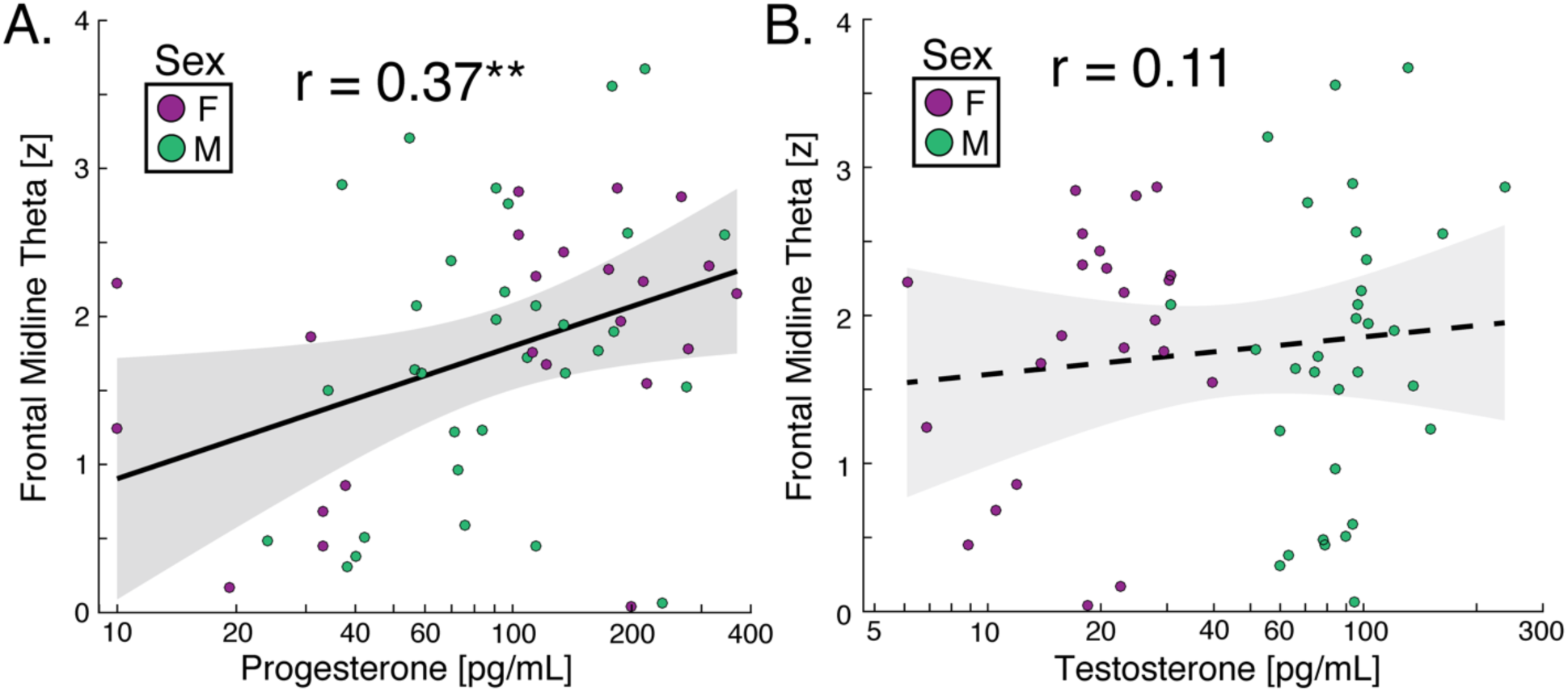
Correlation analysis (Pearson; N=55) of progesterone (A) and testosterone (B) concentration to frontal midline theta oscillatory amplitude. Solid line depicts significant relationship, dotted line is non-significant. Shaded area is 95% confidence interval. ** p < 0.01.

Recent evidence suggests that the aperiodic signal, or background noise, of the human brain is a biologically relevant signal independent of neuronal oscillations (Voytek et al. 2015). With a sharper slope in the background noise, low frequency power is increased, despite no band-specific increase in power. In order to confirm that our results are specific to theta frequency neuronal oscillations and not a result of a difference in background noise, we fit and removed the 1/f aperiodic signal (Figure 2). We found that our initial correlation between FMT amplitude and P4 concentration across all 55 participants (r(54) = 0.377, p = 0.0061) was still significant after 1/f correction (r(54) = 0.284, p = 0.035). There was no correlation between alpha (8-12Hz), beta (15-25Hz), or gamma (25-45Hz) amplitude in FCz and P4 concentration in the raw spectra or after 1/f correction. For comparison, we also analyzed peak theta amplitude in the central-occipital Oz electrode which displays a non-significant and weak correlation with P4 concentration in the raw spectra (r(54) = 0.198, p = 0.148) that was removed by 1/f correction (r(54) = 0.057, p = 0.667). There was no correlation between alpha (8-12Hz), beta (15-25Hz), or gamma (25-45Hz) amplitude in Oz and P4 concentration in the raw spectra or after 1/f correction.

**Figure 2.**
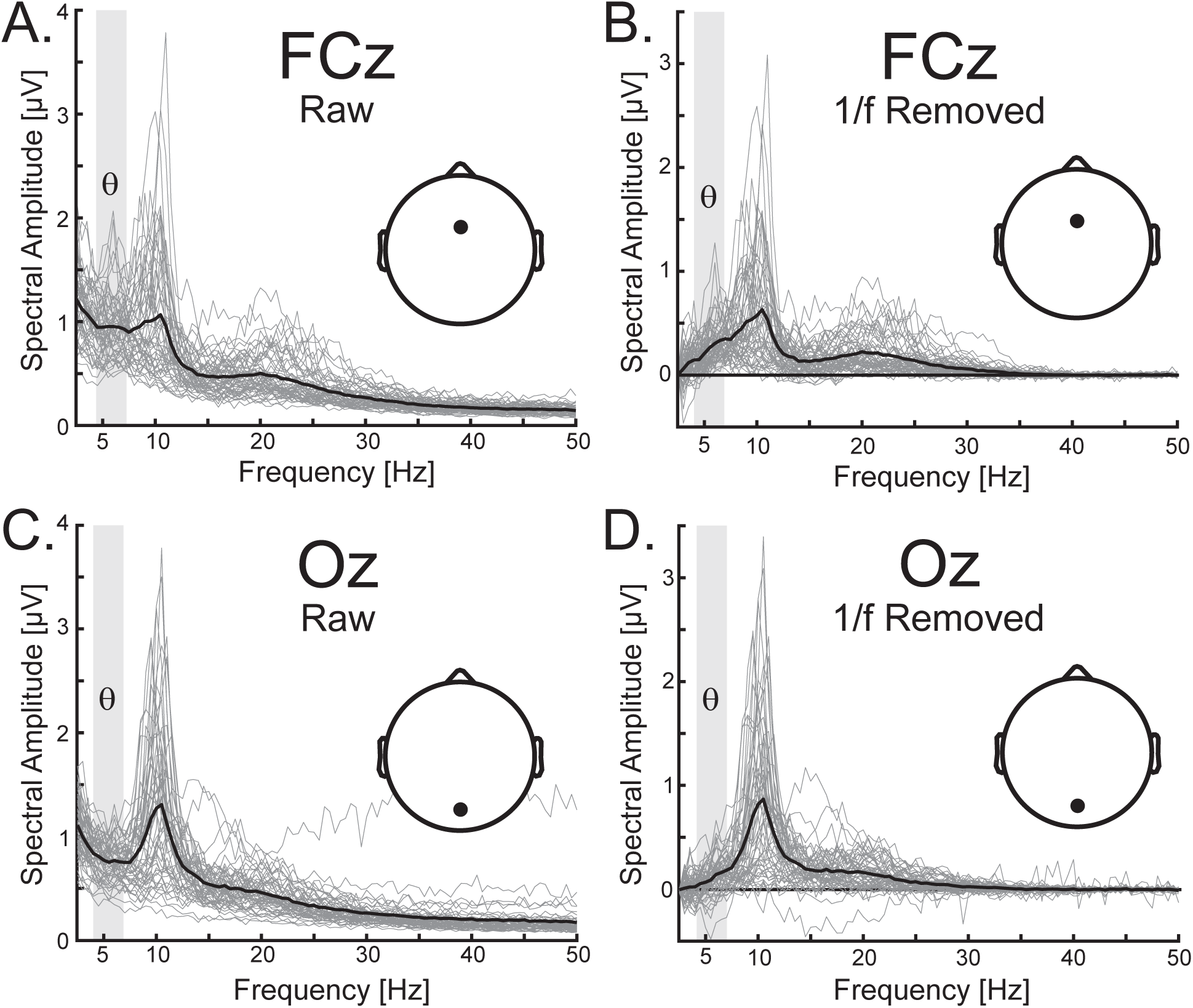
Frontal midline theta oscillations survive aperiodic signal correction. Raw spectral amplitude for (A) FCz and (C) Oz electrode. 1/f correction was a linear regression on the log(amplitude) and log(frequency) values for each participant that was subtracted from the data for (B) FCz and (D) Oz electrode. Individual participants are display in light grey and the group average is displayed in black. Theta band is highlighted in a light grey box. FCz and Oz are depicted on the scalp.

To confirm that the correlation between FMT oscillatory amplitude and P4 concentration is spatially specific to the frontal-midline electrodes, we correlated P4 concentration with theta amplitude for each electrode (Figure 3A). Consistent with the canonical distribution of FMT, the positive correlation with P4 was specific to FCz and its surrounding electrodes. To better understand the neural origin of theta oscillations, we ran source localization on theta oscillations and performed a median split of our participants based on FMT amplitude in sensor-space to analyze the spatial distribution of theta oscillations in source-space between the high and low FMT groups (Figure 3B). As hypothesized FMT oscillations were greatest in bilateral prefrontal cortex and not in medial prefrontal cortex and superior frontal gyrus. These qualitative results for the left hemisphere are displayed in Figure 4A and 4B. We hypothesized that FMT oscillations originated in the frontal-parietal network (FPN) and not the default mode network (DMN) (Yeo et al. 2011). The FPN and DMN are visualized in Figure 4C and 4D to illustrate the spatial consistency between the spatial distribution of theta amplitude in source-space and the independently-derived FPN network.

**Figure 3.**
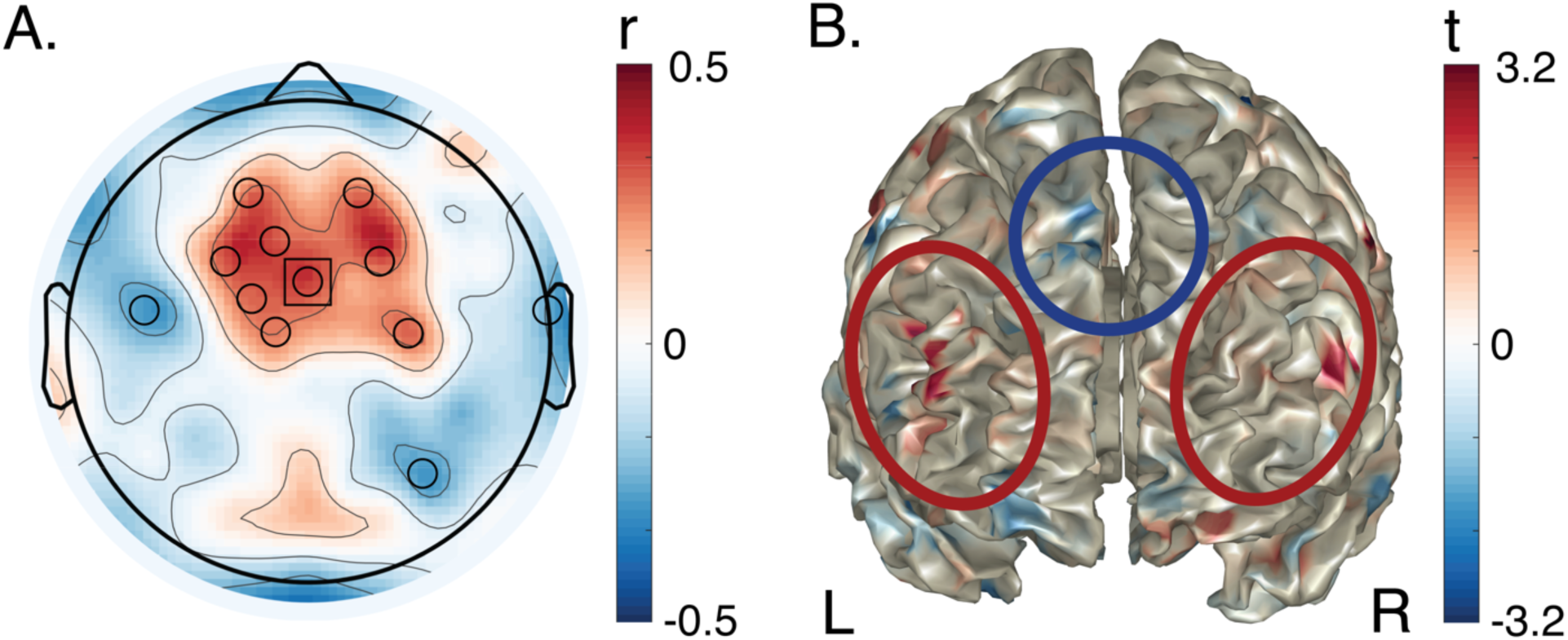
Spatial distribution of frontal midline theta (FMT) oscillations in sensor-space and source-space. (A) Sensor-space correlation (Pearson; N = 55) between theta amplitude and P4 across males and females revealed the canonical distribution of FMT. FCz is outlined with a black square. A black circle depicts a correlation with significance of p < 0.05. (B) Source localization of theta amplitude. For display purposes, voxel-wise independent t-values (df = 53) between participants median split on sensor-space FMT amplitude. Theta oscillations localized to bilateral prefrontal cortex. Anterior-to-posterior axial view of prefrontal cortex. L = left, R = right. The red ellipses highlight increased theta oscillations in the lateral prefrontal cortices, and the blue circle highlights decreased theta oscillations in the superior frontal gyri.

**Figure 4.**
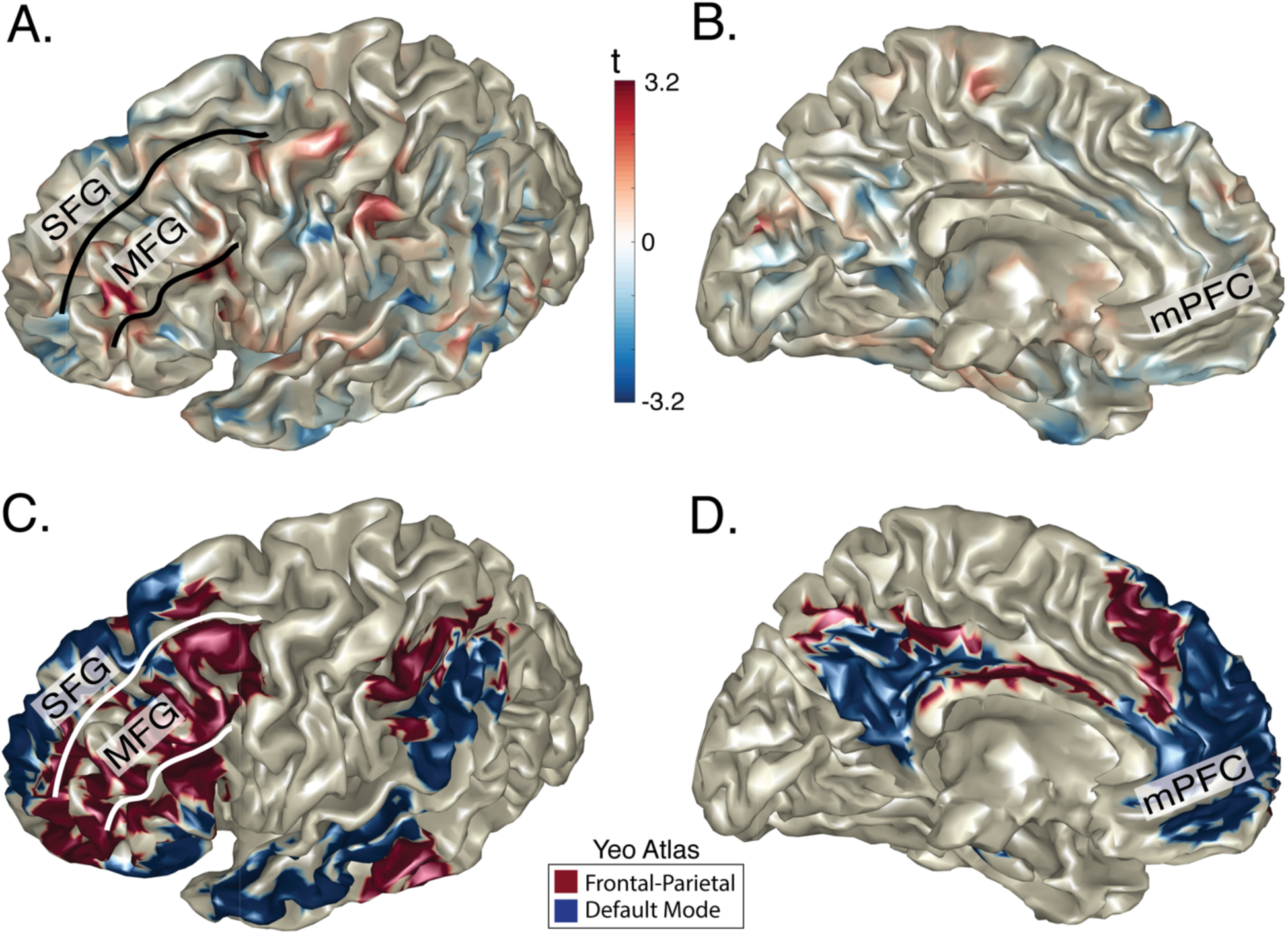
Source localization analysis of frontal midline theta amplitude reveals activation in bilateral middle frontal gyrus (MFG), and not in superior frontal gyrus (SFG) or medial prefrontal cortex (mPFC). Lateral (A) and medial (B) view of the spatial distribution of increased theta amplitude (source-space) with increased frontal-midline theta amplitude (sensor-space) in left hemisphere. Lateral (C) and medial (D) view of the frontal-parietal (red) and default mode (blue) networks from an atlas that was independently derived from resting-state functional connectivity in fMRI (Yeo et al. 2011).

As hypothesized, a one-tailed t-test between participants with high and low FMT found an increase in theta amplitude within the FPN (t(53) = 1.697, p = 0.048, d = .460). There was no increase in theta amplitude within the DMN between the high and low FMT groups (t(53) = - 0.501, p = 0.691, d = .136). Next, we controlled for the increase in theta activity in FPN relative to DMN within-participant and found that theta amplitude was significantly increased in participants with greater FMT amplitude (one-tailed; t(53) = 1.798, p = 0.039, d = .485). Finally, we ran a correlation between theta amplitude derived from source localization and steroid hormone concentrations (Table 2). As expected, we found a significant positive correlation between P4 and theta amplitude in the FPN and in the FPN relative to DMN, but not for the DMN alone. There was no relationship between source localized theta amplitude and T.

**Table 2.** Correlation analysis (Pearson; N = 55) between steroid hormone concentrations and theta amplitude within the frontal parietal network (FPN) and default mode network (DMN). FPN and DMN were independently derived from fMRI analysis. * p < 0.05

|  | Progesterone (P4) | Testosterone (T) |
| --- | --- | --- |
| FPN | **0.343\*** | -0.019 |
| DMN | -0.024 | 0.096 |
| FPN - DMN | **0.310\*** | -0.079 |

FMT is hypothesized to be a cognitive control signal that guides the activity of posterior cortex. One neural mechanism by which low frequency cognitive control signals in prefrontal cortex modulate high frequency bottom-up signals is via phase amplitude coupling. In particular, phase amplitude coupling between FMT phase and parietal-occipital gamma amplitude [30-50Hz] was shown to play a causal role in working memory processes (Berger et al. 2019). This previous experiment found that the phase of FMT coupled to right parietal-occipital gamma oscillations. We ran a phase amplitude coupling analysis between theta phase in the FCz electrode and gamma amplitude in parietal-occipital electrodes (Figure 5). We found that gamma amplitude in right parietal-occipital cortex showed significant coupling with FMT (Figure 5; one-tailed t-test at p < 0.05), and that phase amplitude coupling between FMT and right parietal occipital cortex correlated with P4 concentration (r(54) = 0.288, p = 0.047, one-tailed). This effect was not present in the left occipital-parietal cortex (r(54) = 0.144, p = 0.147, one-tailed).

**Figure 5.**
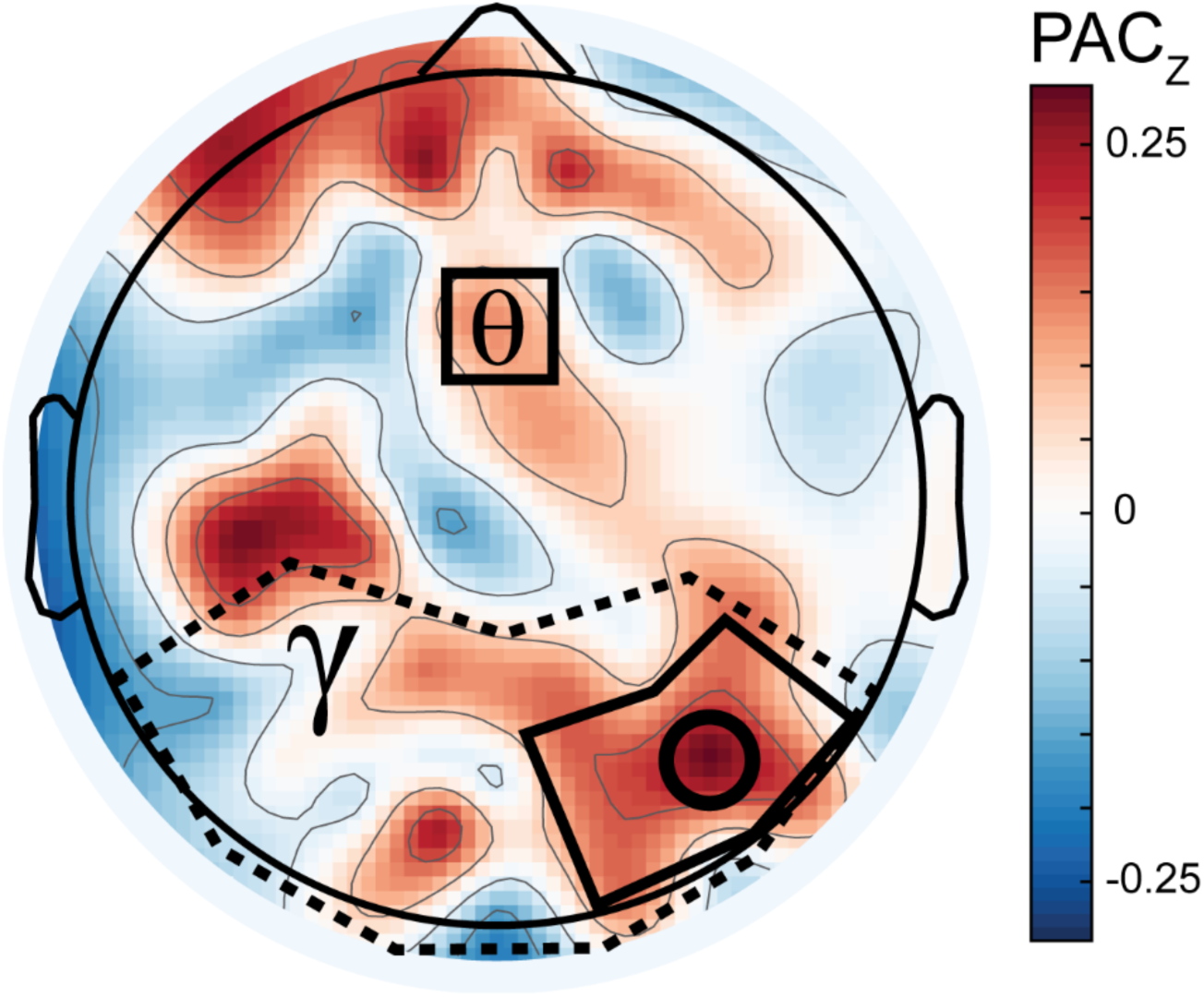
Top-down phase amplitude coupling analysis. Frontal-midline theta phase (FCz electrode, black square) and parietal-occipital gamma amplitude (search light in dashed outline) were significantly coupled in right parietal-occipital cortex using permutation testing (black circle depicts p < 0.05). PAC_Z_ in right parietal-occipital cortex (electrodes for analysis outlined in rectangle) positively correlated with P4 concentrations across participants (Pearson; N = 55).

## Discussion

We investigated if frontal midline theta (FMT) oscillations were related to the concentration of steroid hormones progesterone or testosterone. We found that progesterone (P4), but not testosterone (T), concentrations correlated positively with FMT oscillatory amplitude. We found supporting evidence that theta frequency oscillatory amplitude in frontal midline electrodes in our analysis represented a genuine neuronal oscillation by removing the aperiodic signal, or background spectra, of the brain for each participant. Theta amplitude was source localized to the frontal-parietal network (FPN) and correlated with P4 concentrations. Additionally, we found that the phase of FMT oscillations was coupled to right parietal-occipital gamma amplitude and that this activity correlated with P4 concentrations. Together, these findings suggest that P4 may increase theta oscillations within the FPN that serve as a top-down control signal over sensory-driven posterior cortical activity.

P4 has been found to play a neuroprotective role in both women and men and readily passes through the blood-brain barrier (see (Schumacher et al. 2014) for review). P4, particularly its neuroactive metabolite allopregnanlone (ALLO), is critical to downregulation of hypothalamic-pituitary-adrenal (HPA) activation following stress by facilitating GABAergic inhibitory signaling; and this compensatory response to stress is impaired in depression (Schumacher et al. 2014) (see (Girdler and Klatzkin 2007) for review). ALLO is a potent, positive allosteric modulator of the gamma-aminobutyric acid (GABA_A_) receptor, and it is through this mechanism that P4 exerts antidepressant, antianxiety, and HPA-axis modulatory effects. ALLO was recently approved by the FDA for the treatment of post-partum depression (Osborne et al. 2017; Kanes et al. 2017). Although we did not directly measure ALLO in this study, concentrations of P4 and ALLO are highly correlated (e.g., rs from .80 to .95) in blood and in the CNS (Freeman et al. 1993; Andréen et al. 2005; Palliser et al. 2015). In some cases, such as postpartum, a reduced correlation between P4 and lower concentration of ALLO predicted increased depressive symptoms (Nappi et al. 2001) suggesting an additional mechanism by which P4, a steroid hormone, is converted into its neuroactive metabolite ALLO, despite their strong correlation in most studies with healthy participants.

GABA is the most abundant inhibitory neurotransmitter in mammalian brain (Paredes and Ågmo 1992). GABA_A_ receptors are found throughout the brain, but are found in the greatest concentration in the striatum (Brittain and Brown 2014). The striatum plays a critical role in cognitive control from operant-conditioning (Pagnoni et al. 2002) to decision-making (Balleine, Delgado, and Hikosaka 2007) due to its primary function in inhibiting motor output (Grillner et al. 2005). The FPN and the striatum have reciprocal anatomical projections that contribute to a dynamic hierarchically-organized system for cognitive control (Jarbo and Verstynen 2015; Badre and Nee 2018). A key mechanism by which internal goals are executed is by selective inhibition of distracting percepts (input-gating) and actions (output-gating) via the prefrontal cortex (Munakata et al. 2011; Chatham and Badre 2015). Facilitation of GABAergic signaling by P4 is consistent with an upregulation of inhibitory cognitive control signals in the FPN.

Furthermore, FMT is an emerging biomarker relevant to the treatment of depression. Previous observational studies have found decreased theta frequency amplitude (Saletu, Anderer, and Saletu-Zyhlarz 2010; Pizzagalli, Oakes, and Davidson 2003) and decreased theta frequency functional connectivity (Linkenkaer-Hansen et al. 2005) during the resting-state in patients with major depressive disorder. A recent experiment found that depressed patients that recruited more FMT activity during a working memory task were more likely to respond to non-invasive brain stimulation treatment (Bailey et al. 2018). Variability in recruitment of FMT oscillations during rest also serves as a predictor for pharmaceutical intervention for major depressive disorder (Mulert et al. 2007). Therefore, elucidating contributing factors to FMT may be of critical importance to future interventions.

As in any scientific study, the work presented here has limitations. We only measured peripheral concentrations of P4 and T. Therefore, we cannot draw inferences about concentrations in the brain. However, P4 readily crosses the blood-brain barrier and significantly contributes to central P4 and ALLO concentrations (Hu et al. 2009). Furthermore, we did not collect information on oral contraception, menopausal status, or menstrual phase for the majority of participants. The middle luteal phase of the menstrual cycle is characterized by an acute increase in P4 and the late luteal phase by a decrease in P4. The late luteal phase is associated with mood disturbances such as a irritability and depressed mood (see (Farage, Osborn, and MacLean 2008) for review) and P4 withdrawal contributes to negative mood (Gulinello, Gong, and Smith 2002). A recent review on the effect of progesterone in the menstrual cycle found that P4 was a key predictor of emotional reactivity in women with premenstrual dysphoric disorder (PMDD) (Sundström-Poromaa 2018). P4 concentrations during the late luteal phase positively correlated with increased activity in the amygdala and increased symptoms of anxiety (Gingnell et al. 2014; Gingnell et al. 2012). Consistent with our prediction, previous research using functional neuroimaging found connectivity in the FPN was modulated in the late luteal phase as a function of P4 concentration (Syan et al. 2017). Treatment with ALLO during the luteal phase reduced symptoms of PMDD (Bixo et al. 2017). While speculative, the decrease in P4 during the late luteal phase could downregulate the FPN and decrease cognitive control resulting in hyperactivity of the amygdala in PMDD (Gingnell et al. 2012). Causal investigation using double-blinded administration of P4 or ALLO during a cognitive control task with EEG is required to establish a causal link between P4, FMT, and cognitive control processes.

Finally, non-invasive brain stimulation is a promising new therapeutic that has been used to target the lateral prefrontal cortex in depression (Perera et al. 2016), and can be tailored to target specific brain networks in a frequency specific manner (Alexander et al. 2019; Ahn et al. 2019). In cognitive neuroscience, transcranial alternating current stimulation (tACS) that synchronizes theta oscillations in frontal and parietal regions increases performance during cognitive control tasks (Reinhart and Nguyen 2019; Jaušovec and Jaušovec 2014; Polanía et al. 2012; Vosskuhl, Huster, and Herrmann 2015). Future research should investigate the application of theta frequency tACS targeted to the FPN as a potential treatment for reproductive mood disorders typified by P4 withdrawal.

## Acknowledgements

Thank you to Jhana Parikh, Julianna Prim, and Alec Sheffield for collecting the data, and to Alec Sheffield for preprocessing a subset of the EEG data. This study was funded by the Department of Psychiatry at the University of North Carolina at Chapel Hill and supported in part by the National Institute of Mental Health of the National Institutes of Health under Award Numbers R01MH101547 and R01MH111889 and the postdoctoral training program in reproductive mood disorders T32MH09331502.

